# Re-definition of *claudin-low* as a breast cancer phenotype

**DOI:** 10.1101/756411

**Authors:** Christian Fougner, Helga Bergholtz, Jens Henrik Norum, Therese Sørlie

## Abstract

The claudin-low breast cancer subtype is defined by gene expression characteristics and encompasses a remarkably diverse range of breast tumors. Here, we investigate genomic, transcriptomic, and clinical features of claudin-low breast tumors. We show that *claudin-low* is not simply a subtype analogous to the intrinsic subtypes (basal-like, HER2-enriched, luminal A, luminal B and normal-like) as previously portrayed, but is a complex additional phenotype which may permeate breast tumors of various intrinsic subtypes. Claudin-low tumors were distinguished by low genomic instability, mutational burden and proliferation levels, and high levels of immune and stromal cell infiltration. In other aspects, claudin-low tumors reflected characteristics of their intrinsic subtype. Finally, we have developed an alternative method for identifying claudin-low tumors and thereby uncovered potential weaknesses in the established claudin-low classifier. In sum, these findings elucidate the heterogeneity in claudin-low breast tumors, and substantiate a re-definition of *claudin-low* as a cancer phenotype.

**Contact information:** C.F.

H.B.

J.H.N.

T.S.

## Introduction

The five breast cancer intrinsic subtypes were initially identified by hierarchical clustering of gene expression values of the most stably expressed genes, before and after chemotherapy, in human breast tumors (1, 2). Claudin-low breast tumors did not emerge as an independent group in this analysis. The claudin-low breast cancer subtype was discovered seven years later in an integrated analysis of human and murine mammary tumors (3). The existence of this subtype has later been observed in several independent breast cancer cohorts (4–9), and an analogous claudin-low subtype has been identified in bladder cancer (10, 11).

The claudin-low breast cancer subtype is defined by gene expression characteristics, most prominently: Low expression of cell-cell adhesion genes, high expression of epithelial-mesenchymal transition (EMT) genes, and stem cell-like/less differentiated gene expression patterns (12). Beyond these gene expression features, claudin-low tumors have marked immune and stromal cell infiltration (9, 12), but are in many other aspects remarkably heterogeneous. No specific genomic aberrations accurately delineate claudin-low tumors, and there is a greater variation in mutational burden and degree of copy number aberration (CNA) than in the other breast cancer subtypes (13). Claudin-low tumors are, however, often genomically stable, potentially due to a protective effect of the EMT-related transcription factor ZEB1 (14). Claudin-low breast tumors are reported to be mostly estrogen receptor (ER)-negative, progesterone receptor (PR)-negative, and human epidermal growth factor receptor 2 (HER2)-negative (triple negative), and are associated with poor prognosis (12, 15). The prevalence of claudin-low breast cancer shows striking variability, ranging from 1.5% to 14% of tumors in breast cancer cohorts (5, 7, 8, 12).

An algorithm (predictor) for identifying claudin-low tumors was described with the original characterization of the subtype (12). Briefly, nine claudin-low cell lines were identified by hierarchical clustering of gene expression values of 1906 breast cancer intrinsic genes (16) in a cohort of 52 cell lines. Cell lines were used to build the claudin-low predictor, rather than bulk tumor samples, to minimize immune and stromal infiltration as confounding factors (12). Two centroids were then defined: one for the cell lines with claudin-low gene expression features and one for all other breast cancer cell lines. Claudin-low tumors are identified by correlating a tumor’s gene expression values to the two centroids and defining a tumor as claudin-low if it has stronger correlation to the claudin-low centroid than the other centroid. Importantly, the intrinsic subtypes (basal-like, HER2-enriched, luminal A, luminal B and normal-like) are first identified using the PAM50 predictor (16), and claudin-low subtyping is subsequently performed as an isolated second step (12). In published studies, *claudin-low* is treated as a sixth intrinsic subtype, and the subtype assigned by PAM50 is therefore over-written in claudin-low tumors (5, 8, 9, 12). As a consequence, claudin-low breast tumors have, thus far, been characterized as a single group, without regard for the distribution of the underlying intrinsic subtypes in the given set of claudin-low tumors (8, 9, 12, 13).

In this study, we aim to elucidate the heterogeneity observed in claudin-low breast cancer. By stratifying claudin-low tumors according to intrinsic subtype, we show that the characteristics of claudin-low tumors reflect the intrinsic sub-type to which they are initially assigned. Further, we develop an alternative method for identifying claudin-low tumors, and demonstrate that the nine-cell line claudin-low predictor (12) may be overly inclusive in classifying tumors with marked immune and stromal infiltration as claudin-low.

## Results

### Characteristics of claudin-low breast tumor are delineated by intrinsic subtype

We identified 87 claudin-low tumors (4.6%) in the METABRIC cohort (4, 5) using the nine-cell line claudin-low predictor (12, 17). By intrinsic subtype, the majority of these were classified either as basal-like (51.7%, *n* = 45), normal-like (32.2%, *n* = 28) or luminal A (LumA; 10.3%, *n* = 9) (Fig. 1a, Supplementary Table S1). 14.6% and 15.3% of all basal-like and normal-like tumors, respectively, were identified as claudin-low. All three remaining subtypes were represented in the set of claudin-low tumors, but with a lower prevalence, representing 0.6 - 1.3% of tumors from each sub-type. Only two HER2-enriched and three luminal B (LumB) tumors were classified as claudin-low. These two subtypes were not analyzed further due to low sample numbers.

**Fig. 1.**
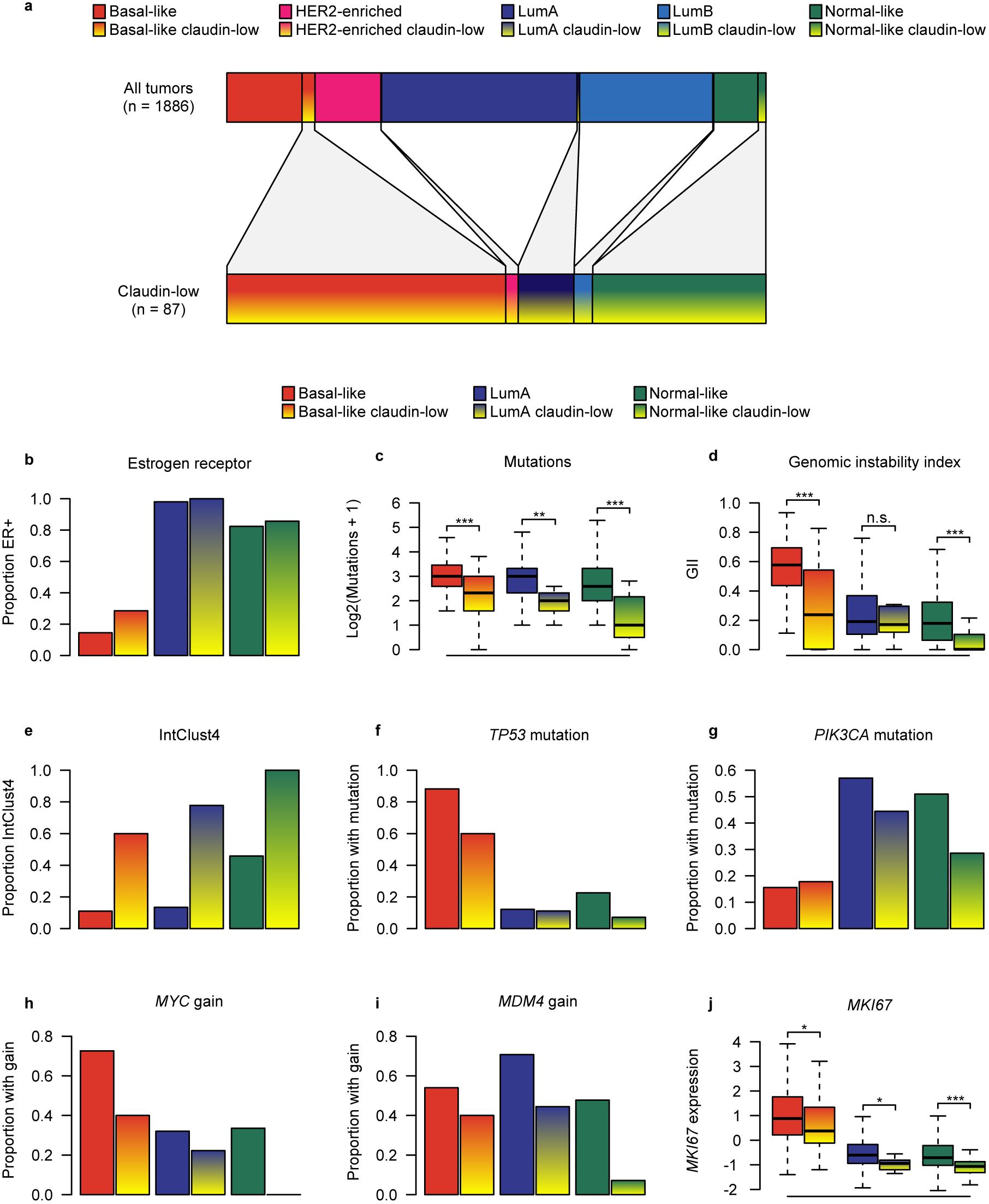
Claudin-low tumors are delineated by intrinsic subtype. **a** Distribution of intrinsic subtypes in the METABRIC cohort for all tumors (top bar, *n* = 1886) and for claudin-low tumors only (bottom bar, *n* = 87). The majority of claudin-low tumors were either basal-like or normal-like. **b** Distribution of estrogen receptor-positivity. 58% of claudin-low tumors were ER-positive. The rate of ER-positivity differed between claudin-low tumors stratified by intrinsic subtype (*P* < 0.001, *χ*^2^ -test). **c** Number of mutations in the panel of 173 sequenced genes. Claudin-low tumors showed lower mutational rates than non-claudin-low tumors of the same subtype. **d** Genomic instability index (GII). Basal-like and normal-like claudin-low tumors showed lower levels of genomic instability than non-claudin-low tumors of the same subtype. **e** IntClust4. 75% of claudin-low tumors were classified as IntClust4. IntClust4 classification differed between claudin-low tumors stratified by intrinsic subtype (*P* < 0.001, *χ*^2^ -test). **f** *TP53* mutation. 38% of claudin-low tumors carried *TP53* mutations. The rate of *TP53* mutation differed between claudin-low tumors stratified by intrinsic subtype (*P* < 0.001, *χ*^2^ -test). **g** *PIK3CA* mutation. 24% of claudin-low tumors carried *PIK3CA* mutations. Differences between claudin-low tumors stratified by intrinsic subtype were not statistically significant (*P* = 0.19, *χ*^2^ -test) **h** *MYC* gain. 26% of claudin-low tumors showed gain of *MYC*. The rate of *MYC* gain differed between claudin-low tumors stratified by intrinsic subtype (*P* < 0.001, *χ*^2^ -test). **i** *MDM4* gain. 30% of claudin-low tumors showed gain of *MDM4*. The rate of *MDM4* gain differed between claudin-low tumors stratified by intrinsic subtype (*P* = 0.006, *χ*^2^ -test). **j** *MKI67* gene expression. Claudin-low tumors consistently expressed lower levels of *MKI67* compared to non-claudin-low counterparts. There were significant differences in *MKI67* gene expression between claudin-low tumors stratified by intrinsic subtype (*P* < 0.001, Kruskal-Wallis test). **All** n.s *P* > 0.05, * *P* < 0.05, ** *P* < 0.01, *** *P* < 0.001. Sample numbers provided in Supplementary Table S1.

There were significant differences in the proportion of tumors expressing estrogen receptor when claudin-low tumors were stratified by intrinsic subtype (Fig 1b; *P* < 0.001, *χ*^2^ -test). 28.6%, 100% and 85.7% of basal-like, LumA, and normal-like claudin-low tumors, respectively, were ER-positive, closely reflecting the pattern seen in non-claudin low tumors (Fig. 1b). These findings indicate that the expression of ER in claudin-low tumors is reflected in their intrinsic subtype, and that characterizing claudin-low tumors as a triple negative subgroup of breast cancer (9, 12) is an over-simplification.

Claudin-low tumors, as a whole, have previously been reported to have a low mutational burden and low level of genomic instability compared to the other subtypes (13, 14). Whole genome copy number data, and sequence data from a panel of 173 cancer-associated genes, was available for the METABRIC cohort (4, 5). When claudin-low tumors were stratified by intrinsic subtype, they consistently showed lower mutational burden and genomic instability compared to their non-claudin-low counterparts (Fig. 1c & 1d), with the exception of genomic instability in LumA tumors. There were, however, also significant differences in mutational burden (*P* = 0.002, Kruskal-Wallis test) and genomic instability (*P* < 0.001, Kruskal-Wallis test) between claudin-low tumors of the different intrinsic subtypes. Despite a degree of subtype specific variations, these findings point toward lower mutational rate and lower levels of genomic instability as *bona fide* claudin-low characteristics.

Curtis *et al.* (4) introduced breast cancer subtypes (IntClust) defined by patterns of CNA with *cis* correlation to gene expression. The genomically stable IntClust4 subtype showed overlap with claudin-low tumors (4). In our analyses, 75% of all claudin-low tumors in the METABRIC cohort were classified as IntClust4. Stratified by intrinsic subtype, claudin-low tumors were consistently more likely to be classified as IntClust4 compared to non-claudin-low tumors of the same subtype (Fig 1e). There were however significant variations in the proportion of claudin-low tumors classified as IntClust4 (*P* < 0.001, *χ*^2^ -test), ranging from 60% of basal-like claudin-low tumors to 100% of normal-like claudin-low tumors. Further, IntClust4 tumors have been separated into ER-positive and ER-negative groups due to major differences in their biological and clinical characteristics, despite strong similarities in gene expression patterns and associated low levels of CNA (4, 5, 18). Claudin-low tumors classified as IntClust4ER+ were predominantly LumA and normal-like, whereas claudin-low tumors classified as IntClust4ER-were predominantly basal-like (Supplementary Fig. S1a & S1b).

The high frequency of claudin-low tumors classified as IntClust4 supports the association between claudin-low gene expression characteristics and genomic stability. However, only 21% of all IntClust4 tumors in the METABRIC cohort were classified as claudin-low, and genomic instability index (GII) did not accurately predict correlation to the claudin-low centroid, as determined by the nine-cell line predictor (12) (Supplementary Fig. S2). Thus, while most claudin-low tumors were genomically stable, only a subset of genomically stable tumors were claudin-low.

No putative driver (19) mutations or CNAs, were found at a significantly higher rate in claudin-low tumors, stratified by intrinsic subtype, than in non-claudin-low tumors of the same subtype (Supplementary Table S2). Rather, claudin-low tumors tended to exhibit patterns of mutation/CNA associated with their intrinsic subtype. Reflecting the lower levels of genomic instability and mutational burden, claudin-low tumors generally had lower incidences of potential driver aberrations compared to their non-claudin-low counterparts. To illustrate the relative frequencies of driver aberrations in claudin-low and non-claudin-low tumors, we selected four early genomic driver aberrations for further analysis: *TP53* mutation, *PIK3CA* mutation, *MYC* gain (located on 8q24), and *MDM4* gain (located on 1q32). Similar to the pattern observed for ER-positivity, the incidence of *TP53* mutations in claudin-low tumors largely followed the incidence seen in the tumors’ intrinsic subtype (Fig. 1f). The differences in *TP53* mutation rates between claudin-low tumors stratified by intrinsic subtype were statistically significant (*P* < 0.001, *χ*^2^ -test). There were similar trends for the other three aberrations analyzed (Fig. 1g - 1i). Claudin-low tumors stratified by intrinsic subtype showed significantly different rates of *MYC* and *MDM4* gain (*P* < 0.001 and *P* = 0.006, *χ*^2^ -test), but not *PIK3CA* mutation (*P* = 0.19, *χ*^2^ -test).

Claudin-low tumors have previously been characterized as slower cycling, with proliferation levels lower than in basal-like tumors, but higher than in LumA and normal-like tumors (8, 12). Ki-67, encoded by the *MKI67* gene, is a commonly used proliferation marker. When claudin-low tumors were stratified by intrinsic subtype, there were significantly different levels of *MKI67* expression between subtypes (Fig 1j; *P* < 0.001, Kruskal-Wallis test), with basal-like claudin-low tumors showing significantly higher levels of *MKI67* expression than LumA claudin-low tumors and normal-like claudin-low tumors (*P* < 0.001 for both, Wilcoxon rank-sum test). Claudin-low tumors did, however, also show significantly lower levels of *MKI67* expression than non-claudin-low counterparts in all intrinsic subtypes (Fig 1j; *P* = 0.01, 0.03 and < 0.001 claudin-low compared to non-claudin-low in basal-like, LumA, and normal-like tumors, respectively, Wilcoxon rank-sum test). Thus, *MKI67* gene expression levels indicate that claudin-low tumors reflect the proliferation levels of their intrinsic subtype but are also slower cycling than non-claudin-low counterparts.

Claudin-low tumors have previously been associated with poor prognosis (8, 12). This characterization was accurate when claudin-low tumors were viewed as a single group (Supplementary Fig. S1c). However, when the survival of patients with claudin-low tumors was stratified by intrinsic sub-type, the survival patterns generally observed in non-claudin-low breast cancer (2) re-emerged (Fig. 2a). Further, there were no significant differences in survival between patients with claudin-low and non-claudin-low tumors within each intrinsic subtype (Fig. 2b - 2d). Thus, we did not find evidence indicating that claudin-low status affects survival in breast cancer patients.

**Fig. 2.**
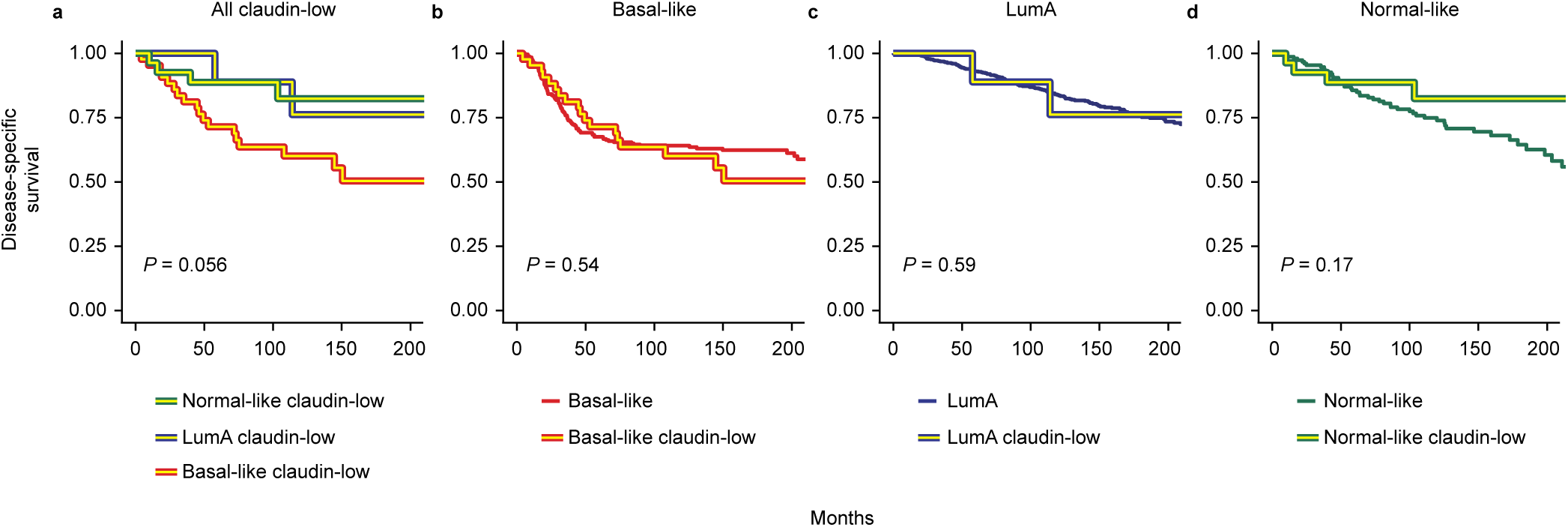
No evidence of claudin-low status as an indicator of poor prognosis in the METABRIC cohort. **a** Disease specific survival in basal-like claudin-low, LumA claudin-low and normal-like claudin-low tumors in the METABRIC cohort. Survival trends recapitulated the patterns seen in non-claudin-low tumors. **b - d**: Disease specific survival in claudin-low and non-claudin-low basal-like (**b**), LumA (**c**) and normal-like (**d**) tumors. Significant differences between claudin-low and non-claudin-low tumors were not found (log-rank test; see sample numbers in Supplementary Table S1).

Claudin-low tumors have been reported to mostly occur in younger patients, with age at diagnosis slightly higher than in basal-like tumors, but lower than in the remaining subtypes (8, 9). When claudin-low tumors were stratified by intrinsic subtype, there were, however, significant differences in the average age at diagnosis (*P* = 0.01, Kruskal-Wallis test; Supplementary Fig. S1d), with basal-like claudin-low tumors diagnosed at a significantly lower age than LumA claudin-low and normal-like claudin-low tumors (*P* = 0.03 and 0.01, respectively, Wilcoxon rank-sum test). There were no statistically significant differences in age at diagnosis between claudin-low and non-claudin-low tumors of the same intrinsic subtype.

### A condensed gene list refines claudin-low classification

Claudin-low tumors have been shown to exhibit high degrees of immune and stromal infiltration (9, 12). Also when stratified by intrinsic subtype, claudin-low tumors in the METABRIC cohort consistently had higher infiltration of immune and stromal cells compared to non-claudin-low tumors (as determined by ESTIMATE, a gene expression-based tool for inferring normal-cell infiltration in tumors (20)) (Supplementary Fig. S1e & S1f). The nine-cell line claudin-low predictor uses 807 genes, and Prat *et al.* acknowledge that it may inappropriately identify some tumors as claudin-low solely due to stromal infiltration (12). We therefore considered whether a more targeted gene list could be used for claudin-low classification, in order to a reduce potentially confounding influence of normal cell infiltration and more accurately isolate features intrinsic to claudin-low tumors.

We created a condensed claudin-low gene list (Supplementary Table S3), consisting of 19 genes representing only the pathognomonic gene expression characteristics of claudin-low tumors: Low expression of cell-cell adhesion genes, high expression of epithelial-mesenchymal transition genes, and gene expression patterns typical of stem cell-like/less differentiated cells (3, 8, 9, 12). In the METABRIC cohort, hierarchical clustering of gene expression values, using the condensed gene list, revealed a tumor cluster with gene expression characteristics in line with those previously described in claudin-low tumors (Fig. 3; *P* = 0.006, SigClust (21)). We refer to tumors in this cluster as *core claudin-low* (*CoreCL*), while claudin-low tumors (as defined by the nine-cell line predictor) outside the CoreCL cluster are referred to as *other claudin-low* (*OtherCL*). Individual inspection of gene expression values showed that OtherCL tumors displayed certain claudin-low characteristics, albeit to a lesser degree than CoreCL tumors (Supplementary Fig. S3).

**Fig. 3.**
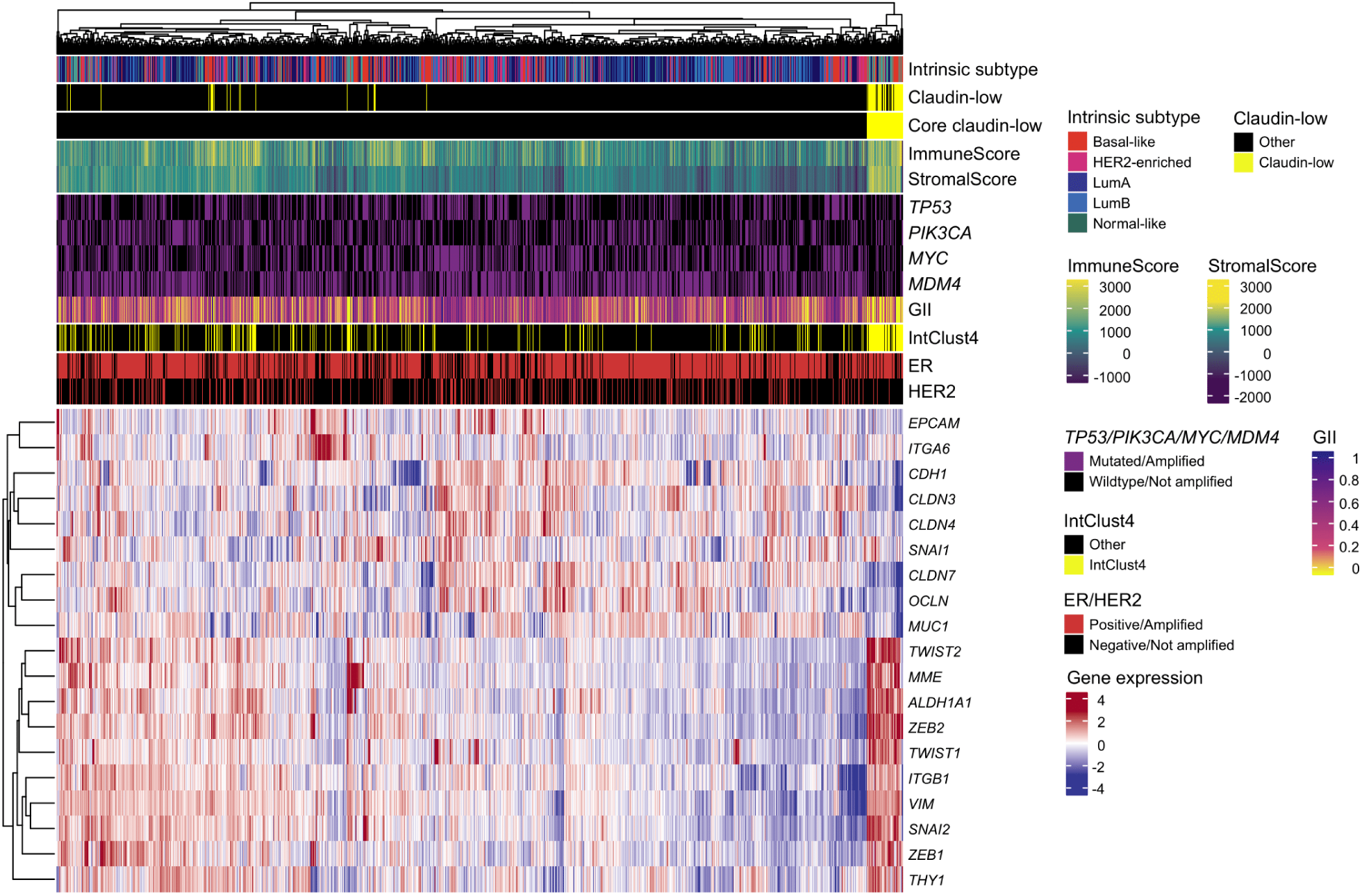
A condensed claudin-low gene list identifies a set of core claudin-low tumors. Heatmap of gene expression values (log2) for a condensed claudin-low gene list in the METABRIC cohort. A cluster, marked *Core claudin-low* (*n* = 79), emerged with transcriptomic and genomic claudin-low characteristics (*P* = 0.006, SigClust (21)).

The CoreCL cluster consisted of 79 tumors (4.2% of tumors in the cohort), of which 57 (72.2%) were identified as claudin-low by the nine-cell line predictor. While several intrinsic subtypes were prominently represented in the group of CoreCL tumors, the OtherCL (*n* = 30) tumors were predominantly basal-like (*n* = 23; Fig. 4a). Thus, our method for identifying claudin-low tumors primarily differed from the nine-cell line predictor by filtering out a set of basal-like tumors with high levels of stromal and immune infiltration (Supplementary Fig. S4a & S4b), but without pathog-nomonic claudin-low gene expression characteristics (Supplementary Fig. S5).

**Fig. 4.**
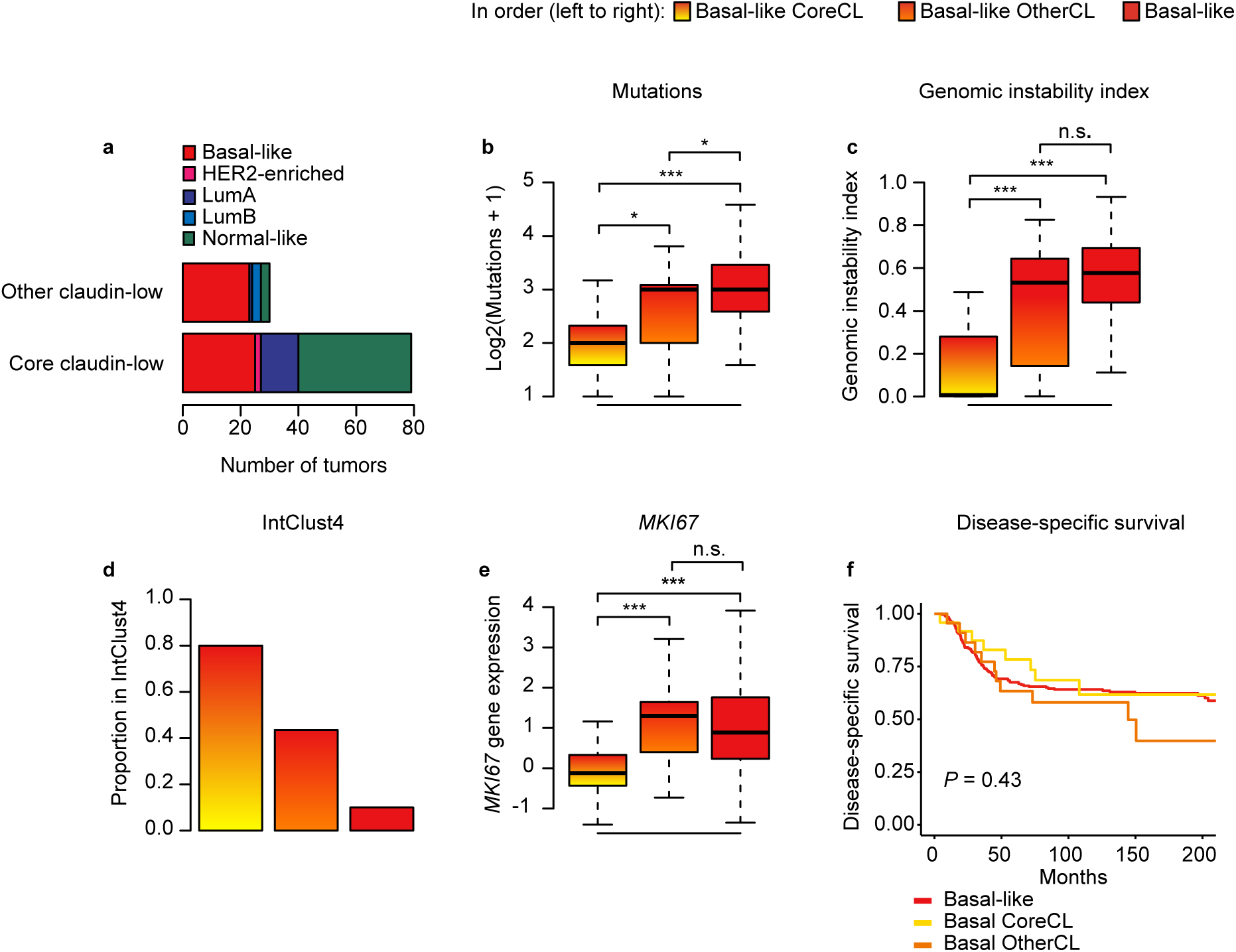
Basal-like OtherCL tumors may be inappropriately classified as claudin-low. **a** Distribution of subtypes in CoreCL and OtherCL tumors in the METABRIC cohort. Hierarchical clustering with the condensed claudin-low gene list filtered out a subset of basal-like claudin-low tumors (as defined by the nine-cell line predictor) with weak claudin-low characteristics. **b** Number of mutated genes in the panel of 173 sequenced genes. Basal-like CoreCL tumors carried significantly fewer mutations than basal-like OtherCL tumors and basal-like non-claudin-low tumors. **c** Distribution of genomic instability index. Basal-like CoreCL tumors showed significantly lower levels of genomic instability than basal-like OtherCL tumors and non-claudin-low basal-like tumors. **d** Proportion of tumors in IntClust4. 80% of basal-like CoreCL tumors were classified as IntClust4. **e** *MKI67* gene expression. Basal-like CoreCL tumors expressed significantly lower levels of *MKI67* than basal-like OtherCL and non-claudin-low tumors. **f** Disease specific survival in basal-like CoreCL, basal-like OtherCL and non-claudin-low basal-like tumors. Disease specific survival in basal-like breast tumors did not significantly differ when stratified by claudin-low status (log-rank test). **All** n.s. *P* > 0.05, * *P* < 0.05, ** *P* < 0.01, *** *P* < 0.001.

There were marked contrasts between the characteristics of basal-like CoreCL tumors (*n* = 25), basal-like OtherCL-tumors (*n* = 23), and non-claudin-low basal-like tumors (*n* = 260). Basal-like CoreCL tumors carried significantly fewer mutations than basal-like OtherCL tumors and non-claudin-low basal-like tumors (Fig. 4b; *P* = 0.015 & *P* < 0.001, respectively, Wilcoxon rank-sum test). Basal-like CoreCL tumors also displayed significantly lower levels of genomic instability than basal-like OtherCL tumors and non-claudin-low basal-like tumors (Fig. 4c; *P* < 0.001 for both, Wilcoxon rank-sum test). There were no significant differences in GII between basal-like OtherCL tumors and non-claudin-low basal-like tumors (Fig. 4c, *P* = 0.082, Wilcoxon rank-sum test). There was also a greater proportion of basal-like CoreCL tumors in IntClust4, than basal-like OtherCL and non-claudin-low basal-like tumors (Fig. 4d, Supplementary Fig. S4c & S4d). In total, 80% of basal-like CoreCL tumors were classified as IntClust4, in contrast to 43% of basal-like OtherCL tumors and 10% of basal-like non-claudin-low tumors. There were also lower rates of *TP53* mutation, *MYC* gain and *MDM4* gain, in basal-like CoreCL tumors compared to basal-like OtherCL and basal-like non-claudin-low tumors, reflecting the lower mutational burden and GII (Supplementary Fig. S4e - S4g). This trend was, however, not evident for *PIK3CA* (Supplementary Fig. S4h). Basal-like CoreCL tumors expressed significantly lower levels of *MKI67* than basal-like OtherCL and basal-like non-claudin-low tumors (Fig 4e; *P* < 0.001 for both, Wilcoxon rank-sum test). There were no significant differences in *MKI67* expression between basal-like OtherCL and basal-like non-claudin-low tumors (*P* = 0.63, Wilcoxon rank-sum test). In sum, the characteristics of basal-like OtherCL tumors show weaker concordance with the characteristics of claudin-low tumors, compared to basal-like CoreCL tumors. The classification of these tumors as claudin-low may therefore be dubious.

Despite differences in genomic and transcriptomic features, as well as in immune and stromal infiltration, there were no significant differences in survival between basal-like CoreCL, basal-like OtherCL and non-claudin-low basal-like tumors (Fig. 4f). These findings reinforce our observations indicating that claudin-low status is not a major determinant of survival in breast cancer patients.

There were few OtherCL samples not classified as basal-like (*n* = 1, 3, and 3 for LumA, LumB and normal-like tumors, respectively; Fig. 4a). The characteristics of normal-like CoreCL (*n* = 39) and LumA CoreCL (*n* = 13) tumors were similar to the characteristics of normal-like claudin-low and LumA claudin-low tumors identified by the nine-cell line predictor (Supplementary Fig. S6). These findings indicate that the nine-cell line predictor identifies certain tumors as claudin-low primarily due to stromal and immune infiltration, and that this may mostly be of concern in basal-like tumors.

### The Oslo2 and TCGA breast cancer cohorts reinforce genomic stability and normal cell infiltration as claud-in-low characteristics

To validate our findings, we queried the Oslo2 cohort (22), for which gene expression data and ER/HER2 status was available. There were 29 claudin-low tumors, as defined by the nine-cell line predictor, in the cohort (7.6%), of which most were classified as basal-like, LumA or normal-like (*n* = 7, 5 and 11, respectively; Fig. S7a). When clustering using the condensed claudin-low gene list, there was a cluster with claudin-low gene expression characteristics and high levels of immune and stromal cell infiltration (Fig. S7b; *P* < 0.001, SigClust (21)). 28 tumors in the cohort (7.3%) were located in the core claudin-low cluster (Fig. S7c), of which 16 (57%) were identified as claudin-low by the nine-cell line predictor. Seven basal-like tumors were classified as claudin-low by the nine-cell line predictor; two of these were CoreCL, both of which were IntClust4, and the remaining five were OtherCL, none of which were IntClust4. Using IntClust4 as a surrogate marker for low levels of genomic instability (4, 18, 23), these findings emphasize that the nine-cell line predictor may be overly inclusive in identifying basal-like tumors as claudin-low. The OtherCL tumors in the Oslo2 cohort were, however, more diverse than in the METABRIC cohort, with 7 of 12 OtherCL tumors being non-basal-like (*n* = 1, 4 and 2 for HER2-enriched, LumA, and LumB, respectively). In total, 89% of CoreCL tumors in the Oslo2 cohort were classified as IntClust4, compared to 38% of OtherCL tumors and 20% of non-claudin-low tumors. Thus, the characteristics of claudin-low tumors in the Oslo2 cohort were mostly consistent with those observed in the METABRIC cohort.

Finally, we explored the TCGA breast cancer cohort (7). 32 of 1082 tumors (3.0%) were classified as claudin-low by the nine cell-line predictor, however, no core claudin-low cluster emerged (Supplementary Fig. S8). As previously noted, normal cell-infiltration is a central characteristic of claudin-low tumors. An inclusion criteria in the TCGA protocol is a tumor cellularity over 60% (7). In comparison, inclusion in the METABRIC cohort requires a tumor cellularity over 40% (4), and there is no such cut-off in the Oslo2 cohort (22). Thus, there may be an association between cellularity cut-off in a cohort and claudin-low prevalence (Fig. 5). This strengthens the observation of normal cell infiltration as a fundamental claudin-low characteristic and may explain the absence of a core-claudin-low cluster in the TCGA-BRCA cohort.

**Fig. 5.**
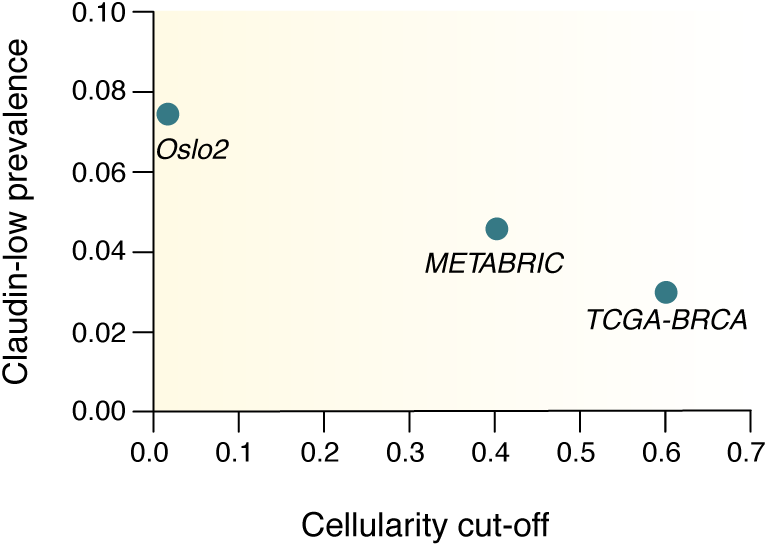
Cut-offs for tumor cellularity may affect the prevalence of claudin-low tumors in breast cancer cohorts. Relationship between cut-offs for tumor cellularity in a cohort and the prevalence of claudin-low tumors.

## Discussion

Here, we have re-evaluated the characteristics of claudin-low breast tumors, from the perspective of *claudin-low* as a phenotype which may permeate the intrinsic subtypes. Through analyses of genomic, transcriptomic and clinical data, we have shown that the characteristics of claudin-low tumors reflect their intrinsic subtype. Characteristics which are attributable to claudin-low status include marked immune and stromal cell infiltration, low levels of genomic instability and mutational burden, and reduced proliferation levels. Finally, we developed an alternative method for identifying claudin-low tumors, and thereby showed that a subset of tumors with pronounced immune and stromal infiltration may be inappropriately classified as claudin-low by the established claudin-low predictor (12).

We stratified claudin-low tumors by intrinsic subtype and found differences between claudin-low tumors of different intrinsic subtypes in almost all aspects analyzed. Perhaps most surprisingly, we found no evidence indicating that claudin-low status affects disease specific survival, contrasting with previous reports of claudin-low as a poor prognosis subtype (8, 12). These findings imply that a large subset of previously reported characteristics of claudin-low tumors are not *bona fide* claudin-low characteristics but are rather an average of the characteristics of several intrinsic subtypes. Thus, the established practice of analyzing claudin-low tumors as a single entity, without taking intrinsic subtype into consideration, may obscure the features that are attributable to claudin-low status.

Claudin-low breast cancer has previously been considered a single disease entity, analogous to the intrinsic breast cancer subtypes (8, 9, 12, 13) (Fig. 6a). Our findings, however, imply that breast tumors are not *claudin-low* instead of the intrinsic subtype to which they are assigned by the PAM50 predictor, rather that they can carry a claudin-low phenotype in addition to their intrinsic subtype (Fig. 6b). According to this interpretation, *claudin-low* is a measure of a set of biological features which is distinct from the set of biological features measured by the intrinsic subtypes.

**Fig. 6.**
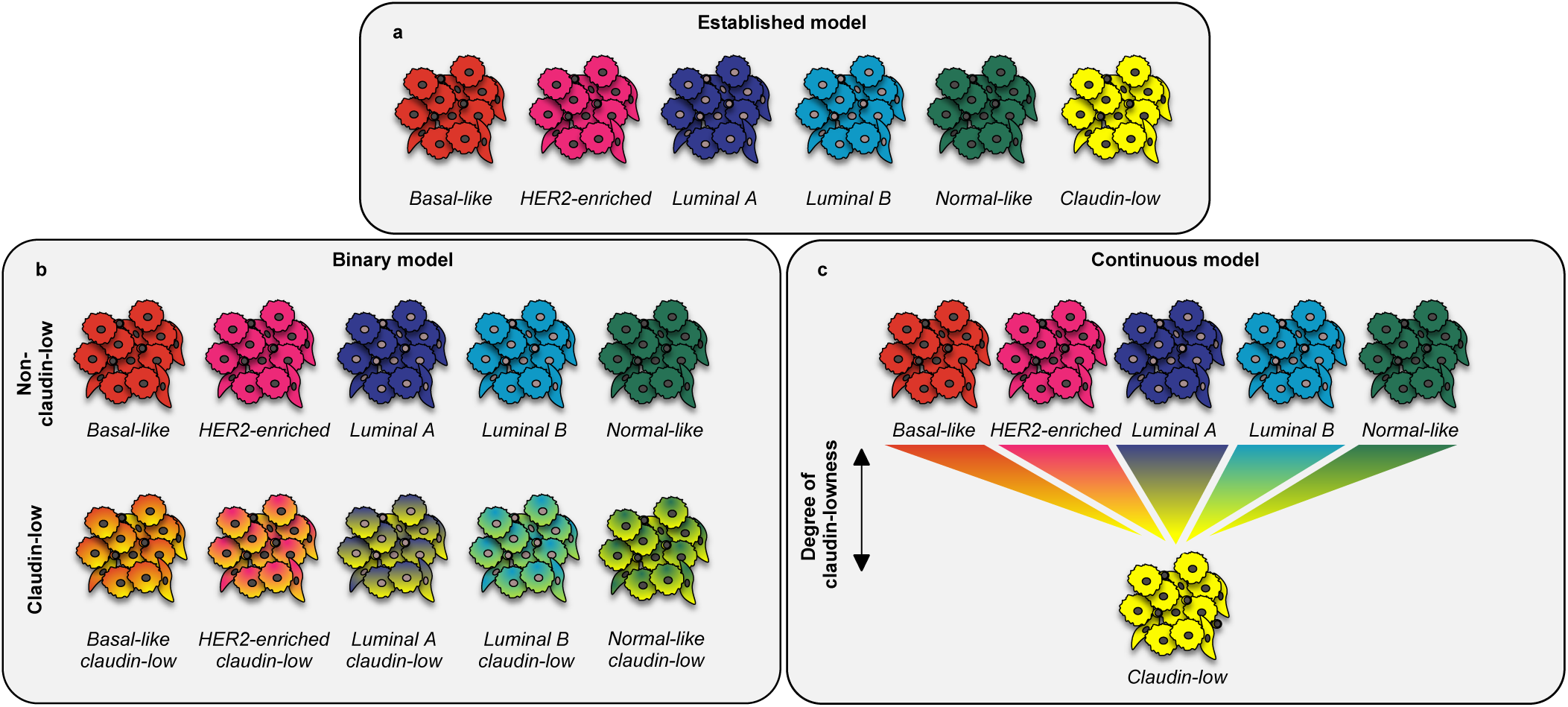
Re-definition of *claudin-low* as a breast cancer phenotype. **a** In the established model, *claudin-low* is interpreted as a sixth subtype, analogous to the intrinsic subtypes. **b** When stratified by intrinsic subtype, claudin-low tumors, however, show characteristics associated with their intrinsic subtype. This implies that tumors are not *claudin-low* instead of their intrinsic subtype, rather that tumors can carry a claudin-low phenotype in addition to their intrinsic subtype. In the binary model, a tumor is either classified as claudin-low, or non-claudin-low. **c** The comparative analysis of CoreCL tumors and claudin-low tumors identified by the nine-cell line predictor, indicates that *claudin-low* may in fact be a continuous feature. Thus, individual tumors may show varying degrees of claudin-lowness, rather than simply being claudin-low or non-claudin-low. In this model, CoreCL tumors, on average, have a higher degree of claudin-lowness than claudin-low tumors identified by the nine-cell line predictor. The continuous model allows for the existence of pure claudin-low tumors, uncoupled from the intrinsic subtypes.

We proposed an alternative method of identifying claudin-low tumors using a condensed gene list. The claudin-low tumors identified using this method (CoreCL) showed more consistent traits than the claudin-low tumors identified by the nine-cell line predictor. OtherCL tumors can be interpreted to not be genuine claudin-low tumors. OtherCL tumors did, however, display some genomic and transcriptomic traits which were consistent with the claudin-low phenotype, though to a lesser degree than CoreCL tumors. A compelling interpretation may instead be that *claudin-low* is a continuum (degree of “claudin-lowness”, Fig. 6c), rather than a binary feature (claudin-low vs. non-claudin-low, Fig. 6b). According to this interpretation, breast tumors exist along a spectrum of claudin-lowness, in which they lie somewhere between: (1) non-claudin-low, fully concordant with an intrinsic subtype, (2) moderately claudin-low with marked imprint of an intrinsic subtype (exemplified by the average claudin-low tumor identified by the nine-cell line predictor), (3) extensively claudin-low, with limited imprint of an intrinsic subtype (exemplified by the average CoreCL tumor), or (4) purely claudin-low, with no imprint of intrinsic subtype (perhaps exemplified by special histological subtypes (24, 25)).

While the continuous model of *claudin-low* may be the most accurate representation of biological reality (26), the interpretation thereof may be challenging for researchers and clinicians, and the binary model may therefore be of greater practical value. Ultimately, a true gold standard definition of *claudin-low* remains elusive, and this will likely be the case until the etiology, pathogenesis and clinical significance of these tumors is precisely understood

Identifying claudin-low tumors using hierarchical clustering of the condensed claudin-low gene list requires a relatively large cohort, and the clustering results are sensitive to the composition of the cohort. The method presented here may therefore not be appropriate for all use cases. An alternative approach to improving the identification of claudin-low tumors may be to only define a tumor as claudin-low if it is identified as such by the nine-cell line predictor and also has a GII under a certain threshold (or is classified as IntClust4). In smaller datasets, it may also be prudent to manually inspect expression levels of genes in the condensed claudin-low gene list in tumors classified as claudin-low by the nine-cell line predictor.

While we did not find evidence that claudin-low status affects survival, certain claudin-low characteristics may nonetheless be clinically relevant and/or actionable. For example, claudin-low tumors show high levels of immune cell infiltration (8, 12), express high levels of PD-L1 (13), are immunosuppressed by T-regulatory cells (27), and carry low mutational burden (13, 14); these factors may all be relevant for the efficacy of immunotherapeutics in claudin-low tumors. The EMT phenotype in claudin-low tumors may in itself be a therapeutic target, and may also have implications for chemoresistance (28). Future studies might address the epigenetic characteristics of claudin-low tumors, and leverage novel single cell sequencing technologies in order to definitively disentangle the features of tumor cells and infiltrating immune and stromal cells. Functional studies will also be necessary in order to elucidate the etiology of claudin-low breast tumors, and to test novel therapeutic modalities. Due to the major influence of normal cell infiltration, it is likely that immunocompetent animal models will be of particular importance for such studies (3, 13, 29, 30).

In summary, we have comprehensively analyzed claudin-low breast tumors, and through these analyses substantiated a re-definition of *claudin-low* as a breast cancer phenotype. We have developed an alternative method for identifying claudin-low tumors and discovered limitations of the established claudin-low predictor. Together, these findings explain a large degree of the heterogeneity observed in claudin-low breast tumors, thereby enabling more accurate and nuanced investigations into this poorly understood form of cancer.

## Methods

### Cohorts

The METABRIC (4, 5), Oslo2 (22) and TCGA-BRCA (7) cohorts were analyzed in this study. Processed data from the METABRIC cohort was downloaded from cBioportal (31, 32); queried data include hormone receptor status, IntClust subtype, disease specific survival, mutation status in a panel of 173 sequenced genes (5), gene-centric copy number status and normalized gene expression values. Intrinsic subtypes (identified using the PAM50 predictor (16)) for the METABRIC cohort were retrieved from supplementary files in Curtis *et al.* (4). Copy number segments and tumor ploidy were retrieved from the repository associated with Pereira *et al.* (5). There were 1886 tumors in the METABRIC cohort with all necessary data.

For the Oslo2 cohort, normalized gene expression values, intrinsic subtypes (identified using PAM50) and hormone receptor status were downloaded from the Gene Expression Omnibus (GEO), accession GSE80999. All 381 samples from the cohort were included in the analyses. Analyses were carried out using GEOquery (33) and Biobase (34). Copy number data was only available for seven claudin-low tumors and was therefore not used in the analyses.

Normalized gene expression values and intrinsic subtypes (identified using PAM50) from tumors in the TCGA-BRCA cohort were downloaded from cBioportal (31, 32). All 1082 tumors from the TCGA-BRCA cohort were analyzed.

### Transcriptomic analyses

The generation and pre-processing of gene expression data is described in the cohorts’ respective publications (4, 5, 7, 22). Gene expression values were mean centered and scaled (z-score). In the Oslo2 cohort, genes represented by multiple probes were reduced to a single gene expression value by finding the mean of all probes representing the given gene.

Claudin-low tumors were identified using the implementation of the nine-cell line claudin-low predictor (12) in the Genefu (17) package for R (35). Euclidean distance was used as the distance metric for claudin-low classification. IntClust subtypes in the Oslo2 and TCGA-BRCA cohorts were determined using a gene-expression based IntClust-classifier (23) implemented in Genefu (17). Immune and stromal infiltration was inferred from gene expression data using *ImmuneScore* and *StromalScore* derived by ESTIMATE (20).

The reduced gene set used to identify core claudin-low tumors (Supplementary Table S3) was manually selected on the basis of published characterizations of claudin-low gene expression features (3, 8, 9, 12). We reasoned that the genes should capture the characteristics unique to claudin-low tumors: Low expression of cell-cell adhesion genes, high expression of EMT genes, and stem-cell like/undifferentiated gene expression pattern. Further, we reasoned that the gene list should not include characteristics which are not unique to claudin-low tumors, such as a low expression of luminal epithelium-related genes. Inclusion of such genes would risk recapitulating the intrinsic subtypes. Hierarchical clustering using the reduced gene list was performed by complete linkage with Euclidean distance as the distance metric. Clustering and visualization was performed using the ComplexHeatmap package (36) for R. The significance of the core claudin-low cluster was evaluated using SigClust (21).

### Genomic analyses

Genomic instability index (GII) was derived by dividing the number of copy number aberrant nucleotides by the total number of nucleotides in the genome. GII was ploidy-corrected by defining a segment as copy number aberrant if the copy number state deviated from the nearest integer value for ploidy. All GII values were ploidy-corrected.

Individually analyzed genomic aberrations were chosen according to the following criteria: (1) known function as early driver events (19, 37); (2) among the most frequently observed aberrations in breast cancer (4, 5); (3) significantly different incidence between intrinsic subtypes (*χ*^2^ -test *P* < 0.05 (4, 5)); (4) non-overlap with other selected events (i.e. only one CNA located on 8q24). *TP53* mutation, *PIK3CA* mutation, *MYC* amplification (8q24), and *MDM4* amplification (1q32) were selected for further analysis on the basis of these criteria.

### Survival Analyses

Survival analyses were performed using the Survival package (38) for R, and Kaplan-Meier plots were generated using the Survminer package.

### Statistical analyses

All significance tests (where applicable) are two-tailed. For continuous variables, Wilcoxon rank-sum test and Kruskal-Wallis test are used to test for differences between two or more than two groups, respectively. For categorical variables, Fisher’s exact test and *χ*^2^ -test are used to test for differences between two or more than two groups, respectively. Significance in survival analyses was determined by log-rank test. Whiskers in box-and-whisker plots are generated using the Tukey method; individual data points are not plotted, as the large number of data points (Supplementary Table S1) and the presence of a small number of extreme outliers tended to obscure overall trends.

## Code availability

All code used in the described analyses is available at https://github.com/clfougner/ClaudinLow.

## Data availability

The data used in this study is available through cBioportal (4, 5, 7, 31, 32), GSE80999 (22), and via supplementary information in Curtis *et al.* (4) and Pereira *et al.* (5). The data used to generate each figure is made available as Source Data.

## ACKNOWLEDGEMENTS

We thank Aleix Prat and Ole Christian Lingj ærde for insightful discussions and critical reading of the manuscript. We are grateful to the Oslo Breast Cancer Research Consortium (OSBREAC) for access to data in the Oslo2 cohort. C.F., H.B. and J.H.N. are supported by grants from the Norwegian Research Council (163027) and South-Eastern Norway Regional Health Authority (2012056) to T.S.

## AUTHOR CONTRIBUTIONS

C.F. conceptualized and designed the study, and performed all bioinformatical analyses. C.F., H.B., J.H.N. and T.S. interpreted the results. J.H.N. and T.S. provided supervision. T.S. acquired funding. C.F. wrote the original manuscript draft. C.F., H.B., J.H.N. and T.S. reviewed and edited the manuscript.

## Supplementary tables

**Table S1.** Distribution of claudin-low tumors by intrinsic subtype in the METABRIC cohort. In total, 87 tumors were classified as claudin-low and 1799 tumors were non-claudin-low.

| Intrinsic subtype | Claudin-low (n) | Non-claudin-low (n) | Proportion claudin-low in subtype |
| --- | --- | --- | --- |
| Basal-like | 45 | 263 | 14.6% |
| HER2E | 2 | 231 | 0.9% |
| LumA | 9 | 684 | 1.3% |
| LumB | 3 | 466 | 0.6% |
| Normal-like | 28 | 155 | 15.3% |

**Table S2. Comparative rates of mutations and CNAs in claudin-low and non-claudin-low tumors (.xslx).** There were no mutations which were found at a significantly higher rate in claudin-low tumors, stratified by intrinsic subtype, than in non-claudin-low tumors of the same subtype. In analyses of core claudin-low tumors (see subheading “A condensed gene list refines claudin-low classification”), OtherCL tumors are treated as non-claudin-low.

**Table S3.**
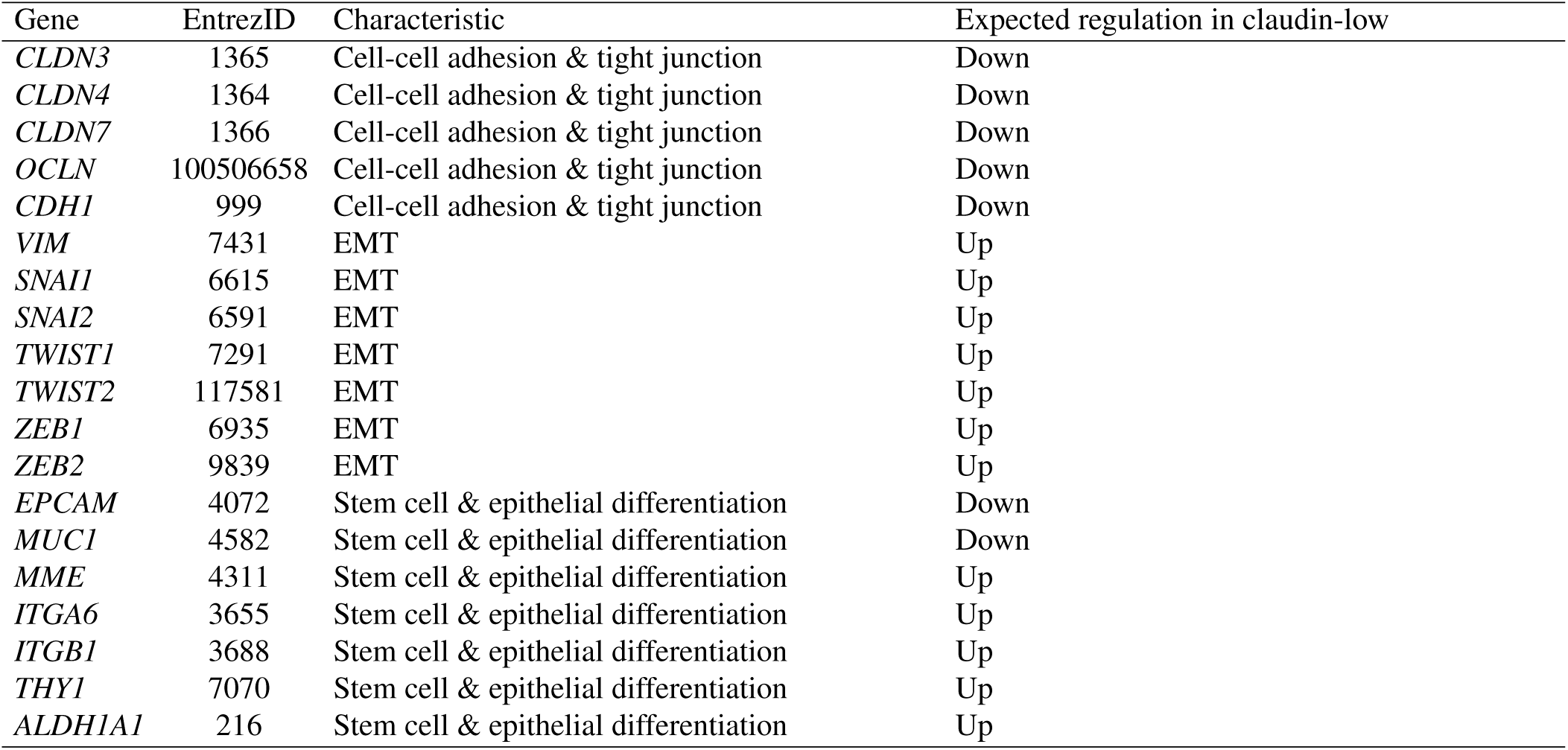
A condensed claudin-low gene list. A list of 19 genes pathognomonic to the claudin-low phenotype. Characteristics/functions listed in the “Characteristic” column are guiding; listed genes may be representative of multiple functions.

## Supplementary Figures

**Fig. S1.**
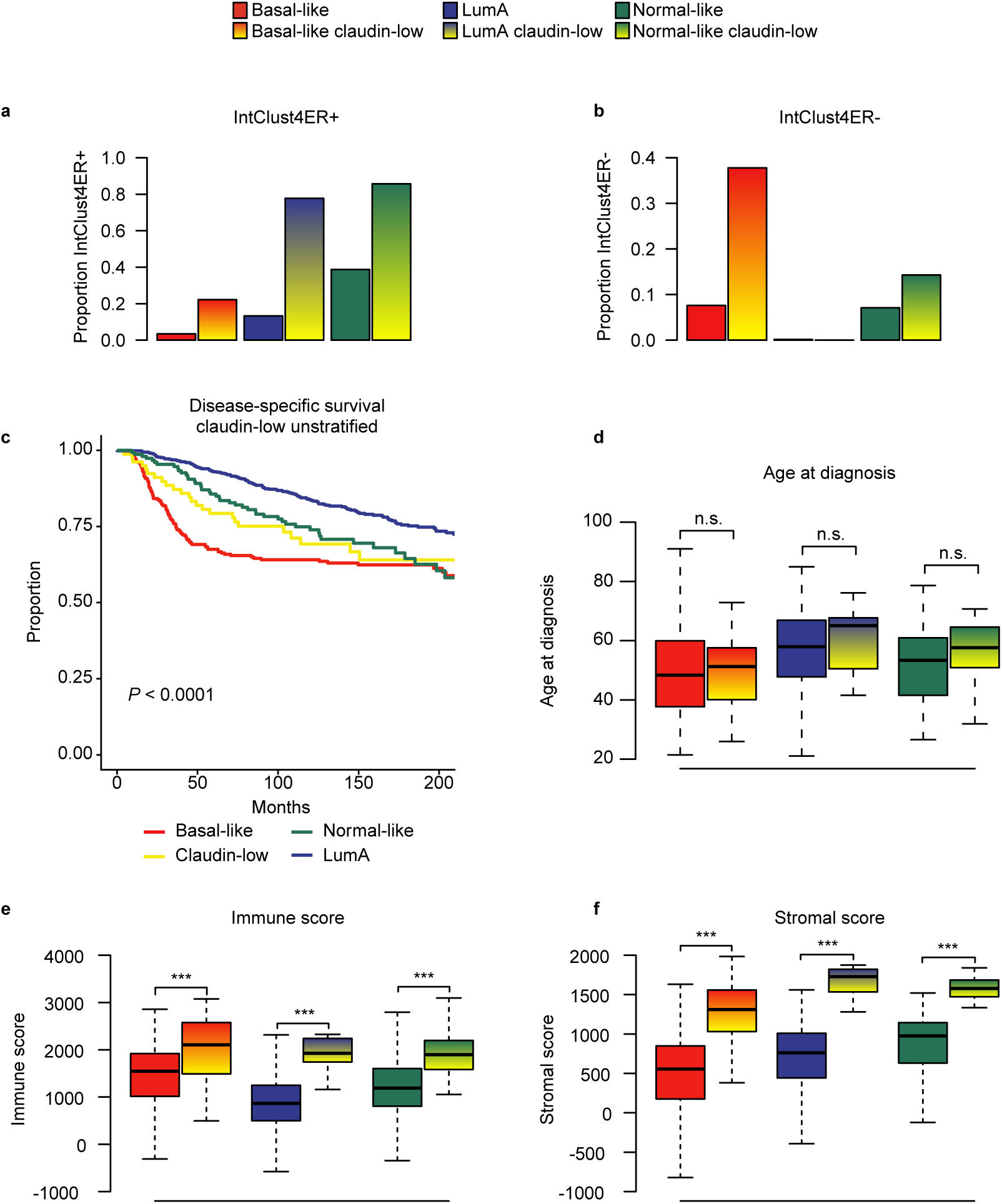
Claudin-low tumors are delineated by intrinsic subtype, continued. **a - b** Proportion of tumors in IntClust4ER+ (**a**) and IntClust4ER- (**b**) by intrinsic subtype and claudin-low status. Basal-like claudin-low tumors tended to be IntClust4ER-, whereas normal-like claudin-low and LumA claudin-low tumors tended to be IntClust4ER+. **c** Disease-specific survival in the METABRIC cohort. Patients with claudin-low tumors are here treated as a single group. The disease-specific survival of patients with claudin-low tumors was superior to that of patients with basal-like tumors, but inferior to that of patients with normal-like and LumA tumors. **d** Age at diagnosis in claudin-low and non-claudin-low tumors. Claudin-low tumors stratified by intrinsic subtype were diagnosed at significantly different ages (*P* = 0.01, Kruskal-Wallis test), with basal-like claudin-low tumors being diagnosed at a significantly lower age than LumA claudin-low and normal-like claudin-low tumors (*P* = 0.01 and 0.03, respectively, Wilcoxon rank-sum test). No significant differences were found between claudin-low and non-claudin-low tumors of the same intrinsic subtype. **e - f** Immune and stromal score in claudin-low and non-claudin-low tumors. Claudin-low tumors of all subtypes had higher levels of immune and stromal infiltration than non-claudin-low tumors of the same subtype. **All** n.s. *P* > 0.05, *** *P* < 0.001.

**Fig. S2.**
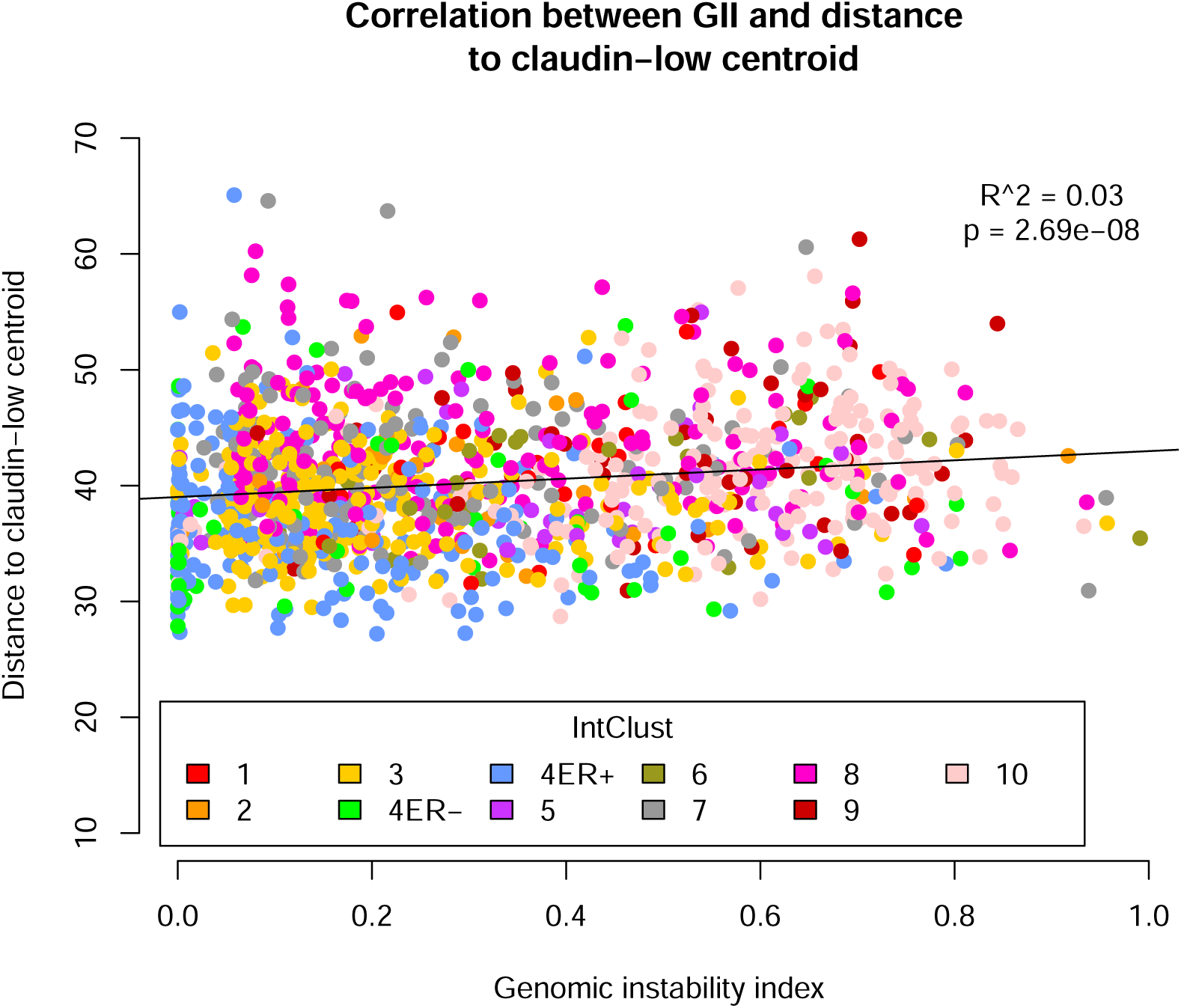
GII alone does not accurately predict correlation to the claudin-low centroid. Correlation (linear regression) between GII and distance to the claudin-low centroid from the nine-cell line claudin-low predictor. While there was a significant correlation between GII and distance to the claudin-low centroid, GII only accounted for 3% of the variance.

**Fig. S3.**
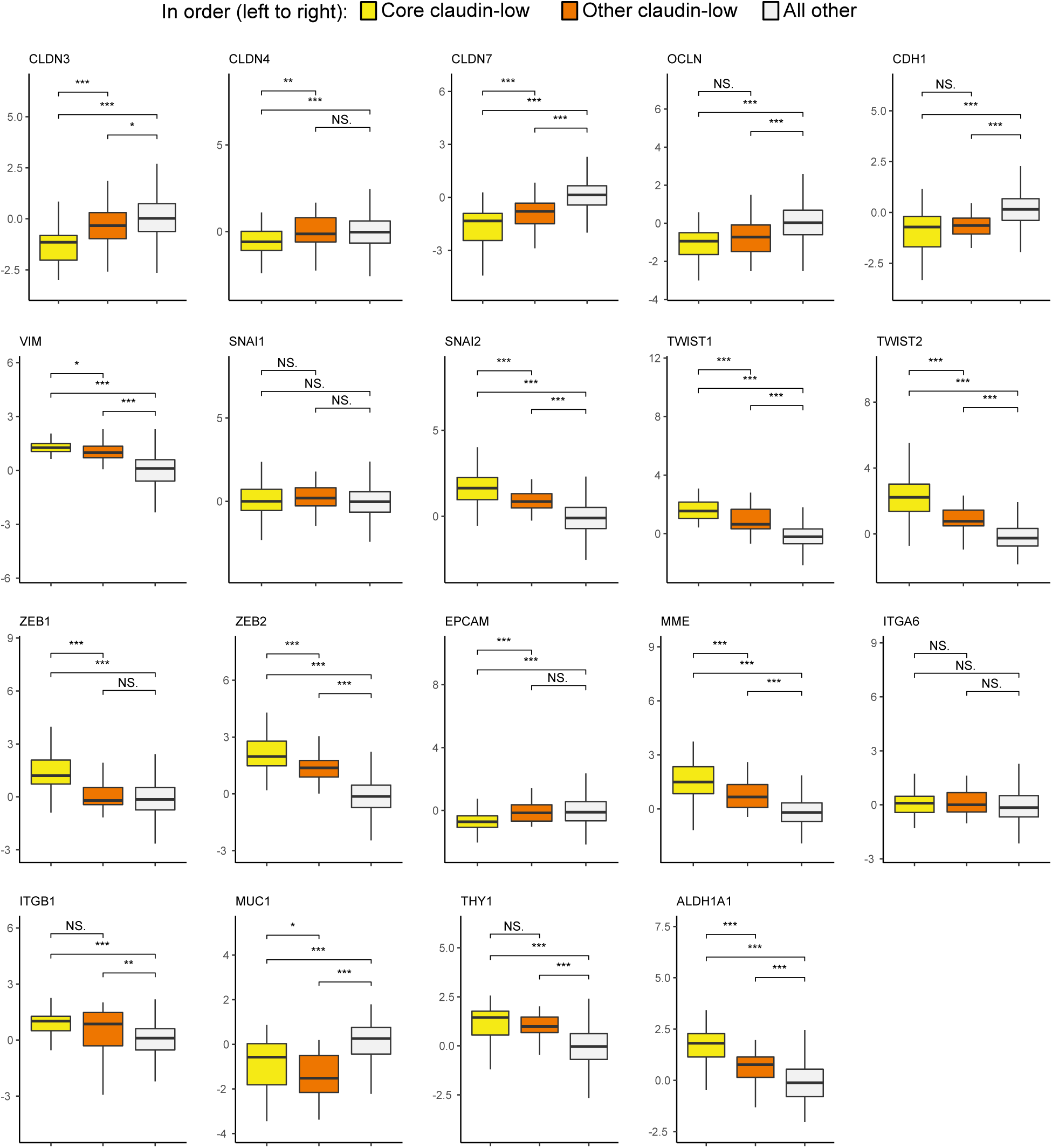
CoreCL tumors show gene expression features consistent with the claudin-low phenotype. Gene expression for all 19 genes in the condensed claudin-low gene list, separated into CoreCL, OtherCL and all other tumors. CoreCL tumors showed gene expression characteristics in line with those previously described for claudin-low tumors. OtherCL tumors showed some claudin-low characteristics, albeit to a lesser degree than CoreCL tumors. NS. *P* > 0.05, * *P* < 0.05, ** *P* < 0.01, *** *P* < 0.001.

**Fig. S4.**
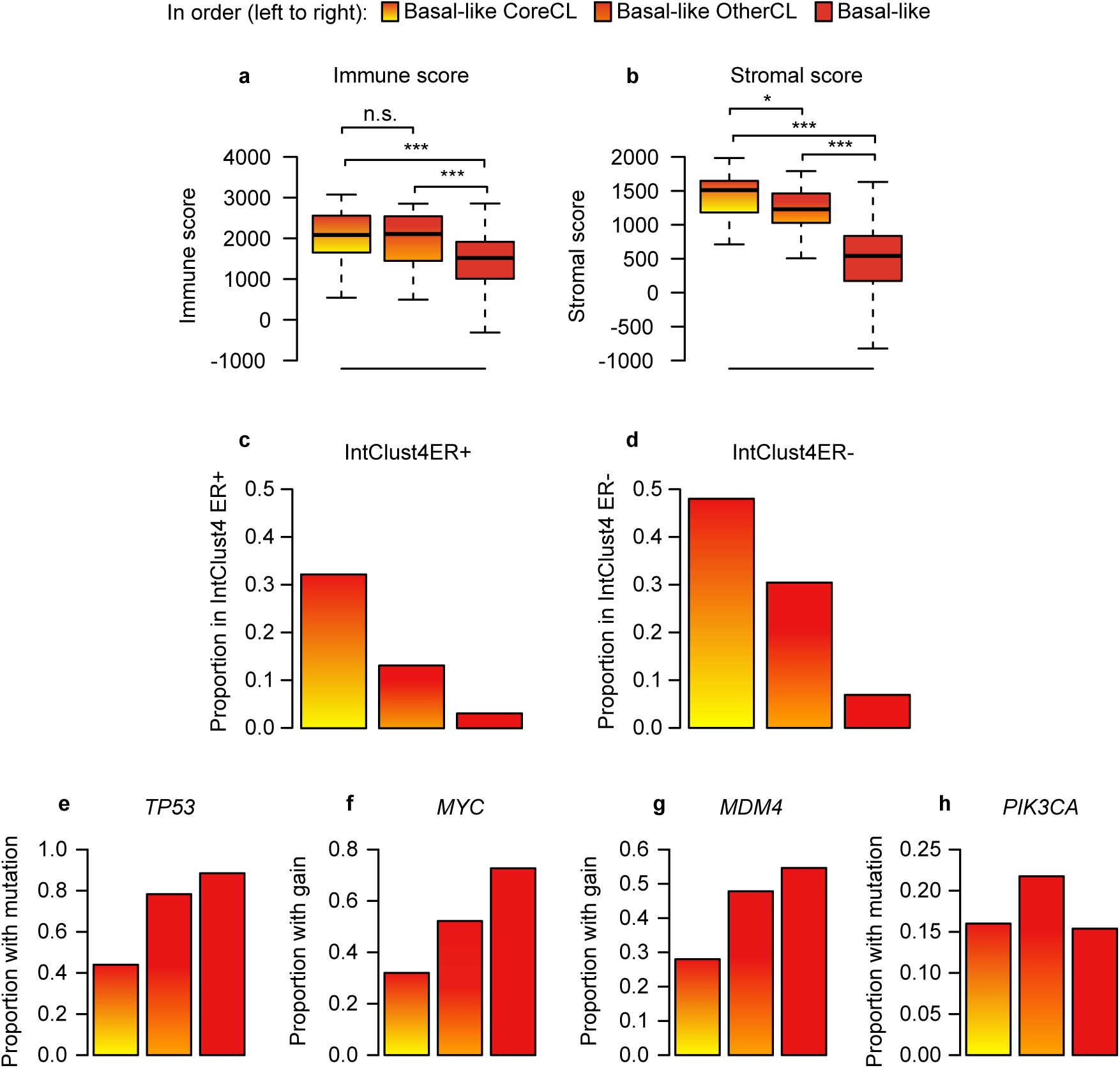
Basal-like OtherCL tumors may be inappropriately classified as claudin-low, continued. **a - b** Immune and stromal score in basal-like core claudin-low tumors, basal-like other claudin-low tumors, and basal-like non-claudin-low tumors. Basal-like CoreCL and OtherCL tumors showed higher immune and stromal infiltration than basal-like non-claudin-low tumors. **c - d** Proportion of basal-like tumors in IntClust4ER+ (**c**) and IntClust4ER- (**d**) by claudin-low status. The majority of basal-like CoreCL tumors were classified as IntClust4, with an overweight of tumors not expressing ER. **e - h** Proportion of tumors with mutation or copy number gain in key genes. Basal-like CoreCL tumors showed lower rates of *TP53* mutation (**e**), *MYC* gain (**f**) and *MDM4* gain (**g**) than basal-like OtherCL and non-claudin-low basal-like tumors. This trend was however not evident in the distribution of *PIK3CA* mutations (**h**). **All** n.s. *P* > 0.05, * *P* < 0.05, *** *P* < 0.00.

**Fig. S5.**
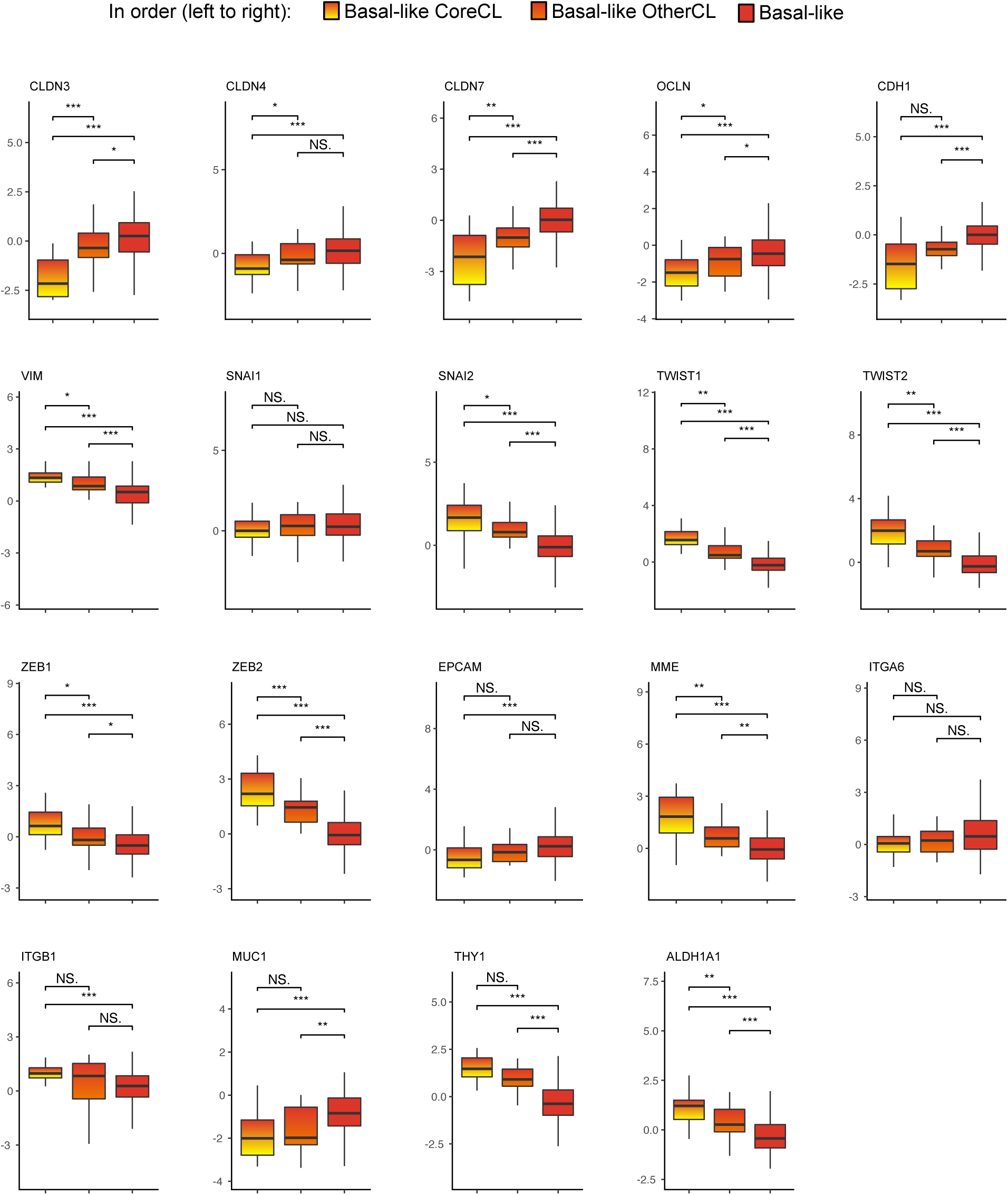
Basal-like CoreCL tumors show claudin-low gene expression characteristics. Gene expression for all 19 genes in the condensed claudin-low gene list for basal-like core claudin-low tumors, basal-like other claudin-low tumors and non-claudin-low basal-like tumors. NS. *P* > 0.05, * *P* < 0.05, ** *P* < 0.01, *** *P* < 0.001.

**Fig. S6.**
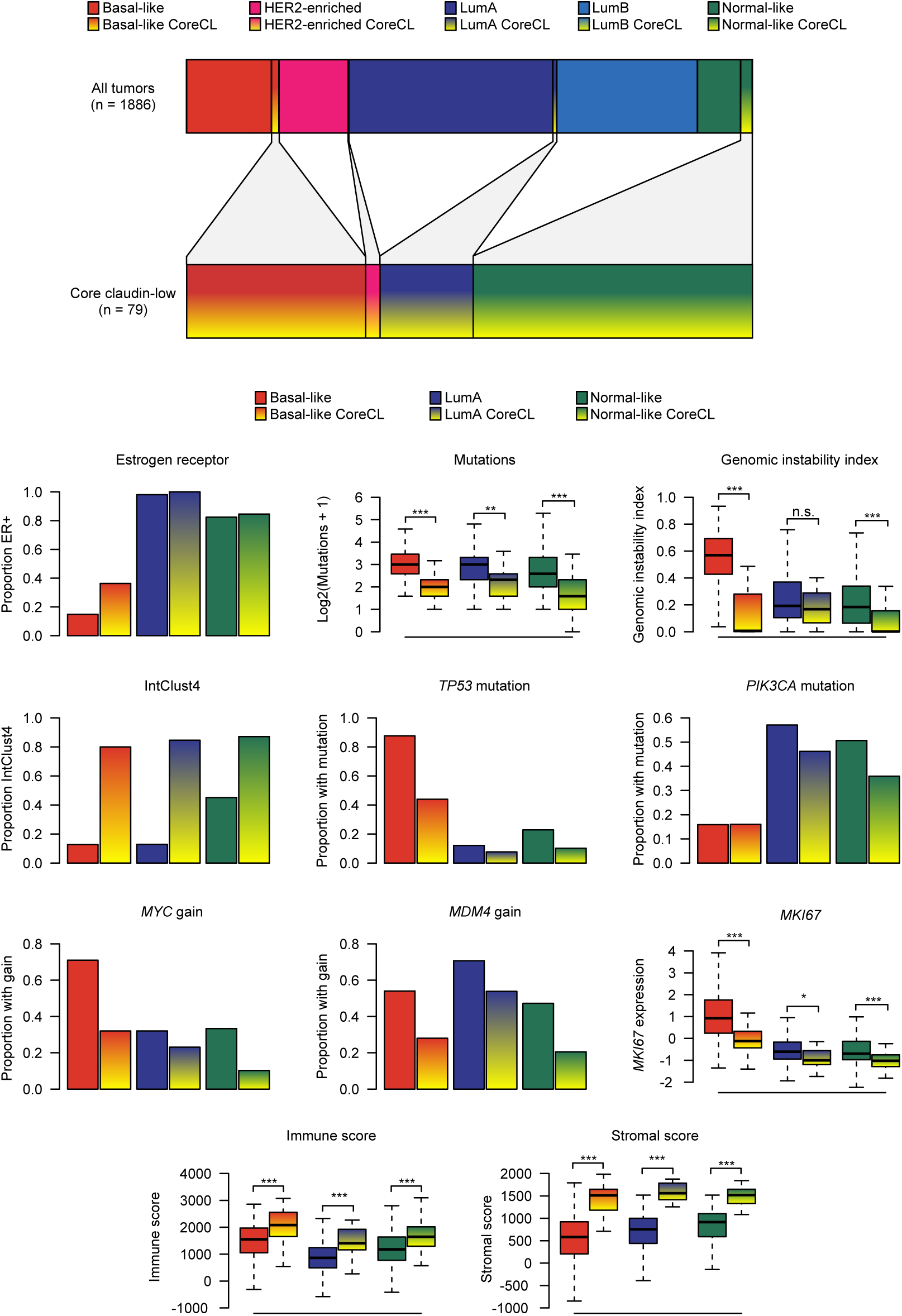
CoreCL tumors are more homogeneous than claudin-low tumors identified by the nine-cell line predictor. Characteristics of CoreCL tumors stratified by intrinsic subtype (panels comparable to Fig. 1 and Supplementary Fig. 1e & 1f). Here, OtherCL tumors are treated as non-claudin-low. There was a reduced variability in the characteristics of basal-like CoreCL tumors compared to basal-like claudin-low tumors as classified by the nine-cell line predictor. The characteristics of LumA CoreCL and normal-like CoreCL tumors were similar to the characteristics of LumA claudin-low and normal-like claudin-low tumors as classified by the nine-cell line predictor. In sum, the characteristics of CoreCL tumors were more homogeneous than the characteristics of claudin-low tumors identified by the nine-cell line predictor. n.s. *P* > 0.05, * *P* < 0.05, ** *P* < 0.01, *** *P* < 0.001.

**Fig. S7.**
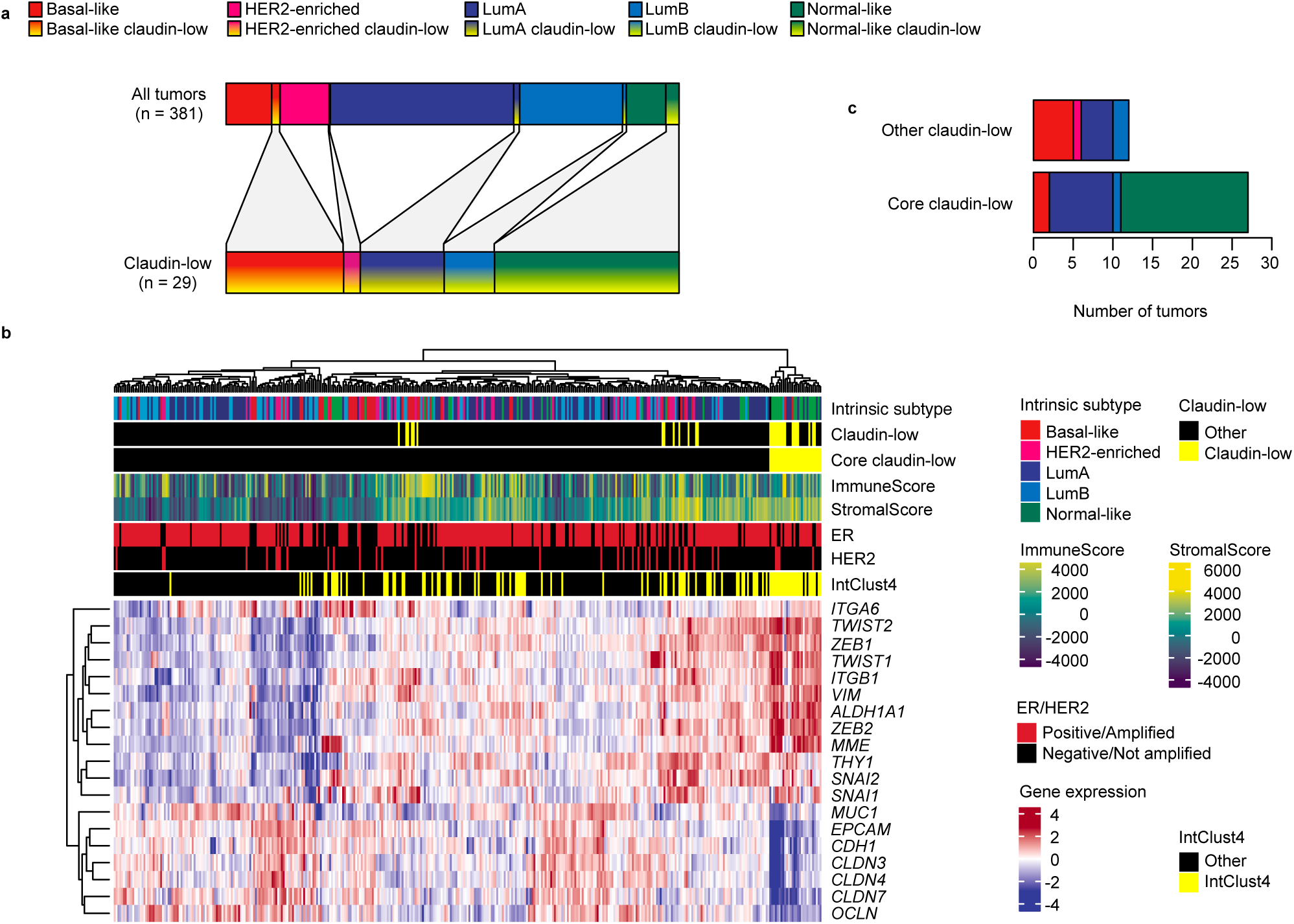
Claudin-low tumors in the Oslo2 cohort recapitulate characteristics observed in the METABRIC cohort. **a** Distribution of intrinsic subtypes in the Oslo2 cohort for all tumors (top bar, *n* = 381) and for claudin-low tumors, as defined by the nine-cell line predictor (bottom bar, *n* = 29). Most claudin-low tumors were either basal-like, LumA, or normal-like. **b** Heatmap of gene expression values (log2) for the condensed claudin-low gene list in the Oslo2 cohort. Hierarchical clustering identified a core claudin-low cluster (*P* < 0.001, SigClust) with similar characteristics to those observed in the METABRIC cohort. Copy number data was not available, however, the representation of IntClust4 in the core claudin-low cluster implies genomic stability in the group. **c** Distribution of subtypes in core and other claudin-low tumors in the Oslo2 cohort. The distribution of subtypes was similar to that seen in the METABRIC cohort, with a slightly larger variation in the intrinsic subtypes of OtherCL tumors.

**Fig. S8.**
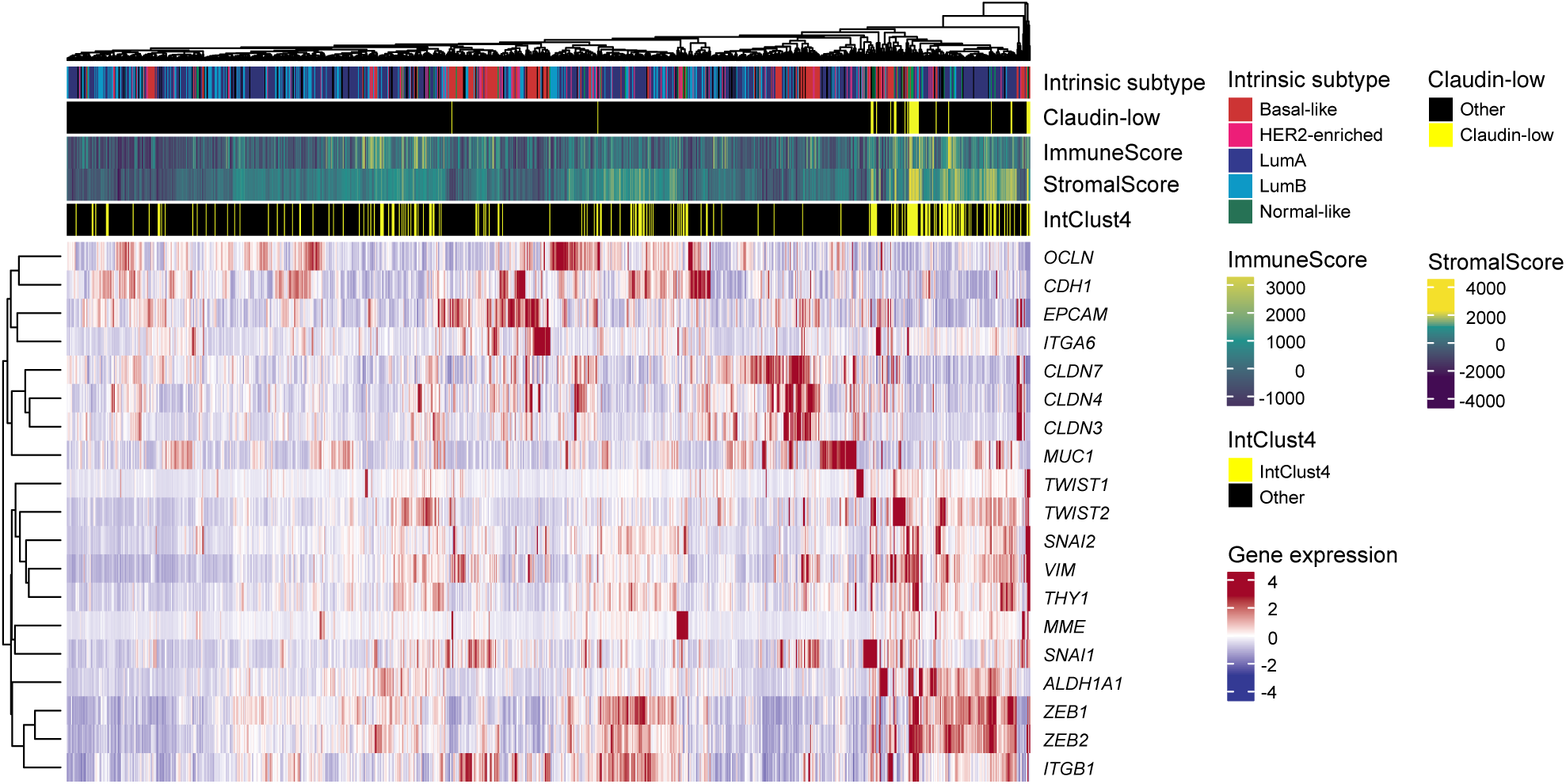
No core claudin-low cluster is evident in the TCGA-BRCA cohort. Heatmap of gene expression values (log2) for the condensed claudin-low gene list in the TCGA-BRCA cohort. No core claudin-low cluster emerged.

